# Loss of the distal-proximal GLP-1/Notch activation gradient in the aging C. elegans germline

**DOI:** 10.1101/2025.10.22.683991

**Authors:** Rustelle Janse van Vuuren, Pier-Olivier Martel, Patrick Narbonne

## Abstract

In *C. elegans hermaphrodites*, the distal tip cells (DTCs) capping the distal gonad arms provide the niche signal, a Notch ligand, to maintain germline stem cell pools. Using fixed germlines, it was recently shown that the transcription of a Notch target gene decreased relatively early-on during adulthood. Here, we used the genetically encoded Notch Sensor Able to detect Lateral Signaling Activity (SALSA) to examine the pattern of GLP-1/Notch activity across the aging distal gonad *in vivo*. Interestingly, we find that the robust and progressively decreasing distal-proximal Notch activation gradient that is observed in young adults gets entirely lost during aging.

## Description

The stem cells’ local microenvironment is essential to maintain their stemness. This microenvironment usually consists of other cells or tissues that immediately surround stem cells and provide them with differentiation-preventing “niche” signals. An example is the activation of a Wnt signal in intestinal stem cells by neighbouring enteroendocrine niche cells, something needed for maintaining their proliferation (Fevr et al., 2007). Although the stem cell microenvironment is known to deteriorate with age, the repercussions within stem cells remain largely unascertained.

In *C. elegans* hermaphrodites, a pool of germline stem cells (GSCs) is located distally within each of two gonad arms. Each GSC pool is maintained by a Notch signal expressed by a single niche cell, the distal tip cell (DTC), which caps the germline and extends processes enwrapping GSCs (Austin and Kimble, 1987; Crittenden et al. 2006). The Notch ligand, LAG-2, present at the surface of the DTC and its processes, activates the GLP-1/Notch receptor on the membranes of adjacent GSCs (Henderson et al., 1997). Upon receptor activation, the Notch intracellular domain (NICD) is cleaved and translocates to the nucleus where it brings a transcriptional activation complex together (Kopan and Ilagan, 2009). This complex activates two main target genes, *lst-1* and *sygl-1*, in a decreasing distal-proximal gradient, to prevent GSC differentiation (Kershner et al., 2014; Lee et al., 2016). As germ cells move proximally due to the ongoing proliferation, they progressively lose Notch stimulation, differentiate and eventually mature into gametes.

A functional decline in germline function occurs with age such that progeny production rapidly decreases in older hermaphrodites, even when sperm supplies are not limited (Hughes et al., 2007). Age-associated alterations in oocyte quality further contribute to the reduction in progeny production (Luo et al., 2009). By the 8^th^ day of adulthood, germline organization is severely disrupted as germline diameter drops and cellular debris and large vacuoles accumulate proximally (Hughes et al., 2011). Part of this functional decline may result from defects in GSC maintenance. Indeed, the progenitor zone (PZ) size progressively decreases with age, going down from 220 cells in young adults to 120 cells in day 5 adults (Kocsisova et al, 2019). A decline in stem cell cycling rates, as governed by reductions in insulin/IGF-1 and MPK-1/ERK signalling levels, may contribute to the decreased PZ size and progeny production (Michaelson et al., 2010; Narbonne et al., 2015; Narbonne et al., 2017; Kocsisova et al, 2019; Robinson-Thiewes et al. 2021). However, the mechanisms underlying the PZ size decline are incompletely understood. Using single-molecule fluorescence *in situ* hybridization (smFISH) targeting a known transcriptional Notch target gene on fixed germlines isolated from aging hermaphrodites, it was established that distal Notch transcriptional activity significantly decreases relatively early-on during adulthood, but not dramatically (Urman et al., 2024). The initially sharp distal-proximal GLP-1/Notch activation gradient nonetheless becomes flatter as the DTCs tend to drift proximally with age, resulting in a larger proximal pool of GSCs having equivalent Notch activity (Urman et al., 2024). The aging worm however undergoes global transcriptional changes, resulting in a decline in transcriptional fidelity (Debès et al., 2023), while mRNA turnover may also be impacted (Borbolis and Syntichaki, 2015). Whether the distal-proximal decreasing gradient of GLP-1/Notch activity persists in older adults *in vivo* remained unclear.

To better characterize the age-related decline in niche signalling, we examined the pattern of GLP-1/Notch activity across the aging distal germline *in vivo*, using the genetically encoded Notch sensor able to detect lateral signalling activity (SALSA) (Shaffer and Greenwald, 2022). The SALSA biosensor consists of a germline-expressed fusion protein made up of GFP, linked via a tobacco-Etch virus protease (TEVp)-cleavable site to a chromatin-tethered RFP (mCherry::H2B). A TEVp is also added to the endogenous Notch intracellular domain (NICD), such that upon receptor activation, NICD::TEVp cleavage and nuclear translocation, GFP is released from the nucleus, resulting in an increased nuclear RFP-to-GFP ratio (Fig. 1A). A strain containing only the fluorescent fusion protein is used as a control to determine the basal RFP/GFP ratio under each condition.

**Figure 1.**
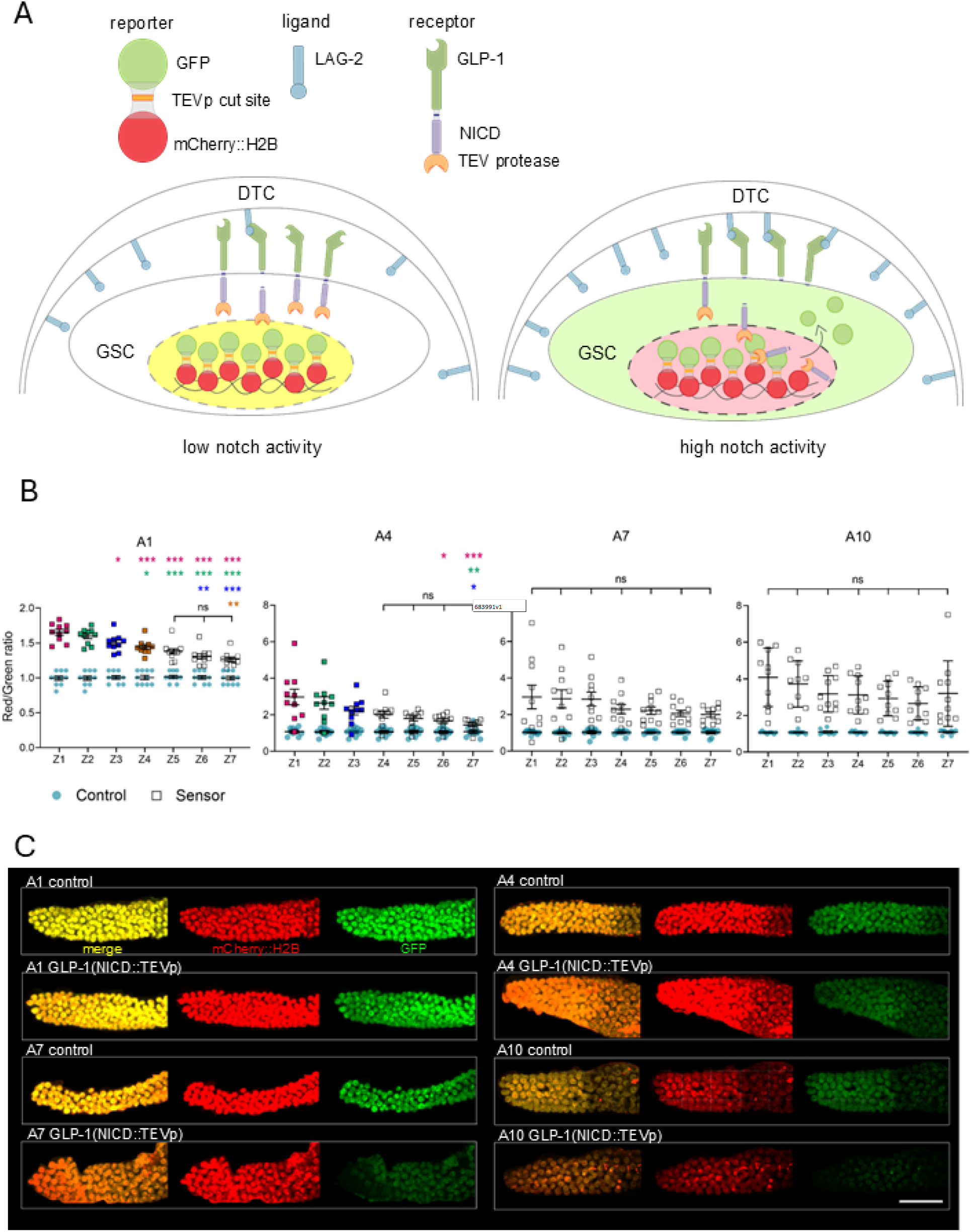
Progressive loss of the spatial GLP-1 activation gradient from the DTC in aging hermaphrodites: **(A)** Schematics of the SALSA biosensor. The sensor consists of a TEV protease (TEVp) fused to the endogenous Notch intracellular domain (NICD), and a separate germline-expressed fusion protein consisting of GFP linked to a chromatin-bound RFP (mCherry::H2B) via a TEVp site. Upon GLP-1/Notch receptor activation, the NICD::TEVp is cleaved, translocates into a GSC nucleus, and cleaves the TEVp site to free GFP from its nuclear RFP::H2B anchor, resulting in an increased nuclear RFP/GFP ratio (Shaffer and Greenwald, 2022). **(B)** Distal gonad nuclear RFP/GFP ratios of adult hermaphrodites were collected every three days from adult day 1 (A1) to 10 (A10) at 20°C, covering nuclei located up to 14 cell diameters away from the DTC, grouped into 7 two-cell diameter zones (Z1-Z7). Sample sizes, 10 gonad arms for each time point. Two independent biological replicates were pooled. Color-coded asterisks mark significance (*p<0.05; **p<0.01; ***p<0.001) versus the corresponding color dataset. The significant decrease in the nuclear RFP:GFP ratio from distal to proximal is lost from A7 onwards. Error bars indicate the standard error of the mean. **(C)** Representative 3D views of distal germlines, constructed from high-resolution confocal z-stacks of immobilized hermaphrodites of the indicated ages and genotypes. Scale bar, 30 μm.

Using this sensor, we measured GLP-1/Notch activation across the PZ, up to 14 cell diameters from the distal tip, in up to 10-day-old adult (A10) hermaphrodites, which corresponds approximately to 2/3 of the typical adult lifespan (Herndon et al., 2002) and is well past the end of the reproductive period (Luo et al., 2009). To this end, germ cells were binned together in seven 2-cell diameter rows from the distal tip, termed zones Z1-Z7. As expected, a robust distal-proximal decreasing gradient of Notch activity was clearly observable in young day 1 adults (Fig. 1B-C). This gradient was however progressively lost during aging as we failed to detect significant differences between zones from day 7 adults onwards (Fig. 1B-C). Thus, GLP-1/Notch receptor activation equalizes across the PZ as animals age.

Although this *in vivo* method allowed to compare Notch activity within living animals of the same age groups, a zone-by-zone quantitative comparison of Notch activity between timepoints was unfortunately not possible. First, nuclear RFP levels varied differentially with age in the sensor and control strains (Figs. 1C and S1). This complication was compounded by a generalized increase in PZ nuclear RFP/GFP ratios over aging in the sensor strain (Fig. 1B-C). For instance, the average nuclear RFP/GFP ratio for Z1 was 1.7 at A1, versus 4.1 at A10 (Fig. 1B). This would counterintuitively suggest that distal GLP-1 activity increases with age. However, this apparent increase in GLP-1 activity in aged GSCs may more likely be a result of their slower turnover, which may allow the NICD::TEVp to release nuclear GFP from distal germ nuclei over a prolonged duration. Moreover, since the ubiquitin–proteasome system becomes dysregulated with age (Koyuncu et al., 2021), the proteolytic NICD::TEVp degradation may become less efficient in aged GSCs. We believe these results highlight important limitations of the SALSA assay towards aging experiments.

Overall, and combined with the literature, our results suggest that an age-dependent reduction in the transcription of direct Notch target genes in GSCs, the proximal drift of the DTC, combined with GSCs’ slower cellular turnover, together effectively lead to the complete loss of the distal-proximal decreasing gradient of Notch signalling activity *in vivo*.

## Methods

### *C. elegans* strains and maintenance

*C. elegans* were maintained at 20°C on standard nematode growth medium (NGM) seeded with *E. coli* bacteria of the strain OP50 (Brenner, 1974). Late L4 hermaphrodites were selected based on vulva development (Seydoux, 1993) and transferred to new plates in batches of 20 until they were collected for analysis, thus for up to 10 additional days. Worms were transferred to new plates on days 2, 5 and 7 to keep their progeny away.

### Imaging

Animals at the indicated ages were paralyzed by soaking in an 8 µL drop of a 0.1% tetramizole (Sigma, L9756) M9 solution on a glass coverslip that was then flipped onto a 3% agarose/M9 pad placed on a microscope slide. Imaging was done within a 30 min window from mounting to minimize any stress signaling (Zellag et al., 2021). The intestine is intertwined with the gonad arms and upon mounting, half of it usually lies on top of one gonad arm. This reduces the detectable fluorescence in that “hidden” arm. Thus, imaging of both gonad arms per animal was rarely done; only when they both lay in the same focal plane and were not obstructed by the intestine. Covered pads were sealed with VALAP (1:1:1 Vaseline, lanolin, paraffin), and 0.35 μm z-step stacks were acquired with a Leica SP8 point scanning confocal microscope and a HC PL APO CS2 40x/1.30 numerical aperture oil objective. GFP and mCherry were excited/collected at 488/495-539nm and 552/590-632nm, respectively, each with 1% laser intensity.

### Quantification and modelling

Notch signalling quantification was performed using Imaris 9.2.1 using the spots tool, by modelling 3.6 mm diameter spheres over individual GSC nuclei based on the red (*mCherry::H2B*) channel. The intensity sums for the red and green channels from within the spheres were measured to generate individual RFP/GFP ratios for each GSC nucleus (Martel et al. *In press*). We considered the cells contained within the first 14 cell diameters of the distal tip, binned into 7 two-cell diameters zones (Z1-Z7). The switchless control RFP/GFP ratios were normalized to 1, to correspond to the expected equimolar nuclear quantities of GFP and RFP; sensor values were transformed according to their respective switchless control average values per zone (Shaffer and Greenwald, 2022).

### Statistical Analysis

Statistical analyses were performed using GraphPad Prism 10.2.0. For multi-group comparisons we used the one-way ANOVA with Tukey’s multiple comparisons when the data set passed the Shapiro-Wilk normality test (Figs. 1B: A1, A7; S1A), and the Kruskal-Wallis with Dunn’s multiple comparisons for sets with non-Gaussian distributions (Fig. 1B: A4, A10). For Fig. S1B, as A1 data failed the Shapiro-Wilk normality test we used again the Kruskall-Wallis test, this time with Mann-Whitney U tests followed by a Bonferroni correction of the resulting P-values.

### Reagents

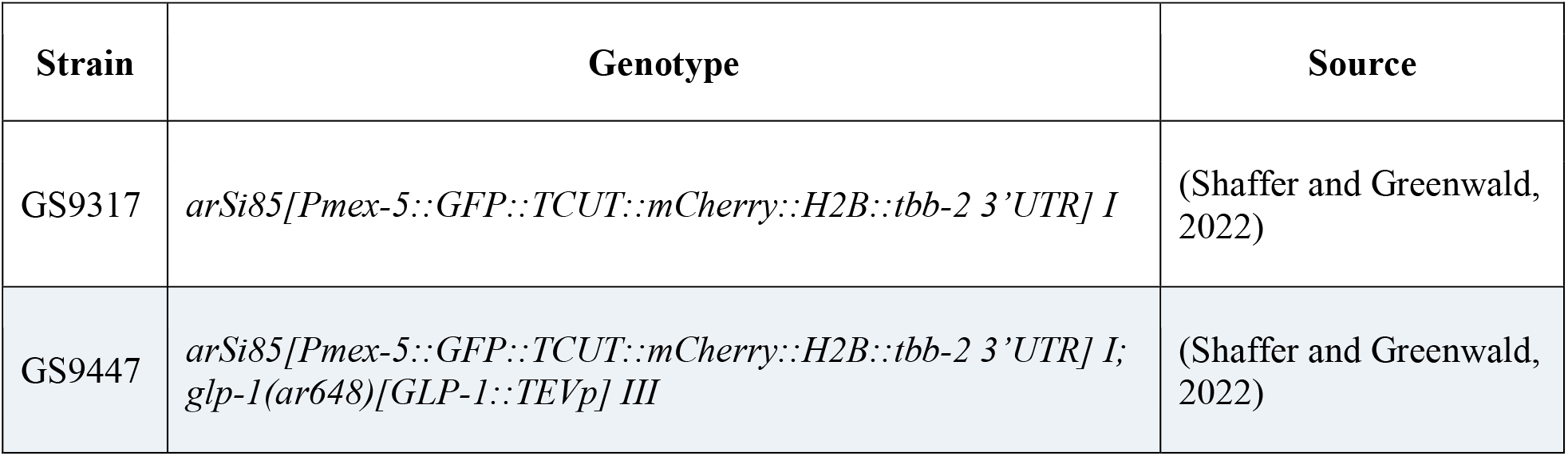
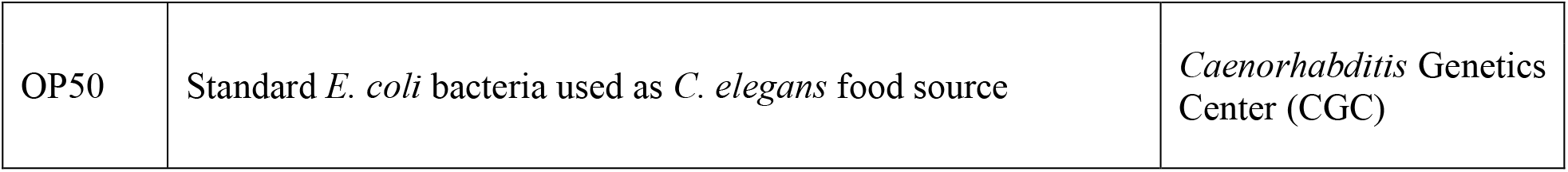

| Strain | Genotype | Source |
| --- | --- | --- |
| GS9317 | *arSi85[Pmex-5::GFP::TCUT::mCherry::H2B::tbb-2 3’UTR] I* | (Shaffer and Greenwald, 2022) |
| GS9447 | *arSi85[Pmex-5::GFP::TCUT::mCherry::H2B::tbb-2 3’UTR] I; glp-1(ar648)[GLP-1::TEVp] III* | (Shaffer and Greenwald, 2022) |
| OP50 | Standard *E. coli* bacteria used as *C. elegans* food source | *Caenorhabditis* Genetics Center (CGC) |

## Acknowledgements

We thank Hugo Germain for sharing his Imaris licence, Iva Greenwald for the SALSA biosensor strains, Alexandre Clouet for verifying statistical analysis as well as WormBase and the CGC for their essential roles in *C. elegans* research. The CGC is funded by NIH Office of Research Infrastructure Programs (P40 OD010440).

## Author Contributions

Rustelle Janse van Vuuren: formal analysis, investigation, data curation, visualization, writing -original draft, conceptualization. Pier-Olivier Martel: conceptualization, methodology, writing - review editing. Patrick Narbonne: funding acquisition, writing - review editing, supervision, conceptualization, formal analysis.

## References

Austin J, Kimble J. 1987. glp-1 is required in the germ line for regulation of the decision between mitosis and meiosis in C. elegans. Cell. 51: 589–599. 1.

Borbolis F, Syntichaki P. 2015. Cytoplasmic mRNA turnover and ageing. Mechanisms of ageing and development. 152:32–42. 2.

Brenner S. 1974. The genetics of Caenorhabditis elegans. Genetics. 77: 71–94. 3.

Crittenden SL, Leonhard KA, Byrd DT, Kimble J. 2006. Cellular analyses of the mitotic region in the Caenorhabditis elegans adult germ line. Molecular biology of the cell. 17: 3051–3061. 4.

Debes C, Papadakis A, Gronke S, Karalay Tain LS, Mizi A, et al., Josipovic N. 2023. Ageing-associated changes in transcriptional elongation influence longevity. Nature. 616: 814–821. 5.

Fevr T, Robine S, Louvard D, Huelsken J. 2007. Wnt/β-catenin is essential for intestinal homeostasis and maintenance of intestinal stem cells. Molecular and cellular biology. 27: 7551–7559. 6.

Henderson ST, Gao D, Christensen S, Kimble J. 1997. Functional domains of LAG-2, a putative signaling ligand for LIN12 and GLP-1 receptors in Caenorhabditis elegans. Molecular biology of the cell. 8: 1751–1762. 7.

Herndon LA, Schmeissner PJ, Dudaronek JM, Brown PA, Listner KM, Sakano Y, et al., Driscoll M. 2002. Stochastic and genetic factors influence tissue-specific decline in ageing C. elegans. Nature. 419: 808–814. 8.

Hughes SE, Evason K, Xiong C, Kornfeld K. 2007. Genetic and pharmacological factors that influence reproductive aging in nematodes. PLoS genetics. 3: e25. 9.

Hughes SE, Huang C, Kornfeld K. 2011. Identification of mutations that delay somatic or reproductive aging of Caenorhabditis elegans. Genetics. 189: 341–356. 10.

Kershner AM, Shin H, Hansen TJ, Kimble J. 2014. Discovery of two GLP-1/Notch target genes that account for the role of GLP-1/Notch signaling in stem cell maintenance. Proceedings of the National Academy of Sciences. 111: 3739–3744. 11.

Kocsisova Z, Kornfeld K, Schedl T. 2019. Rapid population-wide declines in stem cell number and activity duringreproductive aging in C. elegans. Development. 146: dev173195. 12.

Kopan R, Ilagan MXG. 2009. The canonical Notch signaling pathway: unfolding the activation mechanism. Cell. 137:216–233. 13.

Koyuncu S, Loureiro R, Lee HJ, Wagle P, Krueger M, Vilchez D. 2021. Rewiring of the ubiquitinated proteome determines ageing in C. elegans. Nature. 596: 285–290. 14.

Lee C, Sorensen EB, Lynch TR, Kimble J. 2016. C. elegans GLP-1/Notch activates transcription in a probability gradient across the germline stem cell pool. Elife. 5: e18370. 16.

Luo S, Shaw WM, Ashraf J, Murphy CT. 2009. TGF-ß Sma/Mab signaling mutations uncouple reproductive aging from somatic aging. PLoS genetics. 5: e1000789. 15.

Martel PO, Janse van Vuuren R, Degrémont J, Turmel-Couture S, Chaudhari AM, Dologuele E, Beaulieu L, Clouet A. Narbonne P. Niche-associated Type IV collagen promotes GLP-1/Notch receptor activation in the C. elegans germline. Nat. Commun. In press.

Michaelson D, Korta DZ, Capua Y, Hubbard EJA. 2010. Insulin signaling promotes germline proliferation in C. elegans. Development. 137: 671–680. 25.

Narbonne P, Maddox PS, Labbe JC. 2015. DAF-18/PTEN locally antagonizes insulin signalling to couple germline stem cell proliferation to oocyte needs in C. elegans. Development. 142: 4230–4241. 17.

Narbonne P, Maddox PS, Labbe JC. 2017. DAF-18/PTEN signals through AAK-1/AMPK to inhibit MPK- 1/MAPK in feedback control of germline stem cell proliferation. PLoS genetics. 13: e1006738. 18.

Robinson Thiewes S, Dufour B, Martel PO, Lechasseur X, Brou AAD, Roy V, et al., Narbonne P. 2021. Non-autonomous regulation of germline stem cell proliferation by somatic MPK-1/MAPK activity in C. elegans. Cell reports. 35 19.

Seydoux G, Salvage C, Greenwald I. 1993. Isolation and characterization of mutations causing abnormal eversion of the vulva in Caenorhabditis elegans. Developmental biology. 157: 423–436. 20.

Shaffer JM, Greenwald I. 2022. SALSA, a genetically encoded biosensor for spatiotemporal quantification of Notch signal transduction in vivo. Developmental cell. 57: 930–944. e6. 21.

Urman MA, John NS, Jung T, Lee C. 2024. Aging disrupts spatiotemporal regulation of germline stem cells and niche integrity. Biology Open. 13: bio060261. 22.

Zellag RM, Zhao Y, Poupart V, Singh R, Labbe JC, Gerhold AR. 2021. CentTracker: a trainable, machinelearning–based tool for large-scale analyses of Caenorhabditis elegans germline stem cell mitosis. Molecular Biology of the Cell. 32915–930. 23.

