## Supplemental Figure 1 for "Loss of the distal-proximal GLP-1/Notch activation gradient in the aging C. elegans germline"

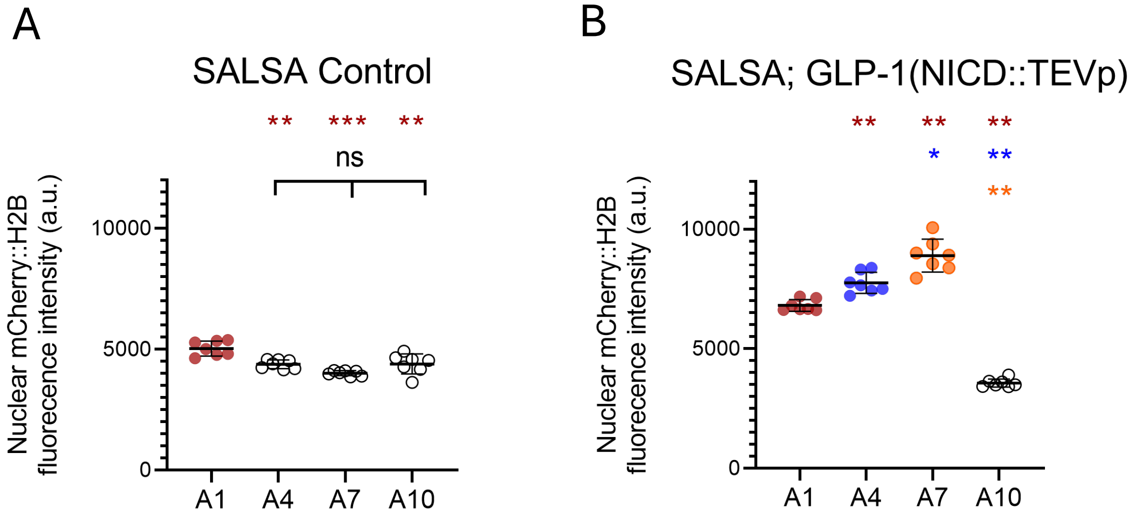


**Figure S1: Inconsistent variations in nuclear mCherry::H2B levels between the GLP-1(NICD::TEVp and SALSA control aging stains. (A-B)** The nuclear mCherry::H2B fluorescence sum intensity was evaluated in the distal gonad arms of adult hermaphrodites of the indicated ages, at 20°C, covering nuclei located up to 14 cell diameters away from the DTC, divided in two cell diameter zones (Z1-Z7). Each data point represents the average sum intensity of nuclear mCherry::H2B for one of these seven zones as averaged from 10 distal gonad arms. Two independent biological replicates were pooled. Color-coded asterisks mark significance versus the corresponding color dataset. *p<0.05; **p<0.01; ***p<0.001.
